# Argon inhibits autophagy and improves cerebral ischemia-reperfusion injury via PI3K/Akt/mTOR pathway

**DOI:** 10.1101/2025.10.31.685964

**Authors:** Shuang Ma, Minghui Wang, Zhongshan Luo, Philip Petersen, Endi Wang, Yu-Xin Xia, Lian-He Yang

## Abstract

It has been reported that argon has neuroprotective effects during cerebral ischemia-reperfusion injury (CIRI). However, the specific mechanism is not fully understood. The aim of this study is to investigate if argon could regulate autophagy to exert a neuroprotective function. A temporary middle cerebral artery ischemia mouse model (tMCAO, ischemia 3h, reperfusion 1h) and the HT22 cell line Oxygen glucose deprivation reperfusion (OGDR) model were used for argon treatment. Transcriptome sequencing and bioinformatics analysis were performed on HT22 OGDR models. Western blot, CCK-8, LDH, transmission electron microscopy, agonists and inhibitors of autophagy and PI3K/Akt/mTOR were used to investigate the specific mechanism. Argon could reduce infarct volume, improve weight recovery, and neurological scores. Enrichment analysis of the KEGG signaling pathway was highly correlated with the PI3K/Akt pathway. Western blot, CCK-8, LDH, and transmission electron microscopy showed that argon could activate the PI3K/Akt/mTOR pathway and inhibit autophagy, promoting a neuroprotective function. After the combined use of autophagy inducer 3, Akt inhibitor MK2206, and mTOR inhibitor rapamycin, respectively, the neuroprotective function of argon was significantly reversed. Our study suggests that argon activating the PI3K/Akt/mTOR pathway and inhibiting autophagy is an important mechanism for its neuroprotective function.

## **1.** Introduction

CIRI refers to the significant damage caused to brain cells when the cerebral blood supply is interrupted and then restored, which commonly leads to a bad prognosis of stroke. It is a complex pathological process involving multiple cellular and molecular mechanisms[1–9]. Currently, the treatment methods and efficacy for CIRI are limited[1, 2, 4, 7–9]. Intra-arterial thrombolysi and mechanical thrombectomy are widely used in clinical practice nowadays, so cerebral ischemia-reperfusion injury (CIRI) is very common. Therefore, choosing an effective and safe method to inhibit cerebral ischemia-reperfusion injury is very important.

In recent years, a series of studies have shown that argon may have neuroprotective functions[10–13]. Argon gas is non-toxic and cheap, although most studies have shown that argon has neuroprotective effects on ischemic brain injury, there is still controversy over whether argon has neuroprotective effects on CIRI[14, 15].On the other hand, the mechanism studies of argon neuroprotective function are very scattered, the current main research findings suggest that argon therapy may have anti-apoptotic properties, which may be related to the involvement of argon in regulating extracellular signal-regulated kinase (ERK)-1/2 [16], heme-oxygenase-1 [17], nuclear factor (erythroid derived 2)-like 2 (Nrf2) [18], TLR2, and TLR4 related signaling pathways[19].

Studies show an increased number of autophagosomes in animal CIRI model brain tissue and the level of autophagy-related proteins, which indicates that cerebral CIRI can activate autophagy. It has been reported that autophagy is moderately activated under conditions such as ischemia and hypoxia, which not only protects cells in nutrient deficient states but also promotes cell survival. However, autophagy is a “double-edged sword,” as excessive activation of autophagy during CIRI can lead to cell lysis and promote cell death, known as autophagic cell death. This autophagic cell death is often accompanied by apoptosis and necrosis, and inhibiting this excessive autophagy may be an effective method to improve CIRI [20–26].

In previous study, we found that argon treatment can improve the neurological score and reduce infarct volume in a rat model of permanent middle cerebral artery obstruction (MCAO) model [27]. In this study, we treated the transient middle cerebral artery occlusion (tMCAO) mouse model and the mouse HT22 cell line OGDR model with argon, then performed transcriptome sequencing and bioinformatics analysis. We found that argon inhibits the autophagy pathway and exerts neuroprotective effects during CIRI via activating the PI3K/Akt/mTOR pathway.

## 2. Materials and Methods

### 2.1. Establishment of mouse tMACO model, argon treatment and neurological score

Specific pathogen-free (SPF) animal feeding was used. Kunming mice were used in this experiment (Liaoning Changsheng Biotechnology Co., Ltd). Only males were selected, with a weight range of 20-25 grams (a total of 68 mice). Mice were divided into the Sham surgery N_2_ group (70% N_2_/30% O_2_, n=15, referred to as Sham N_2_ group), Sham surgery argon group (70% argon/30% O_2_, n=15, referred to as Sham argon group ), tMCAO surgery N_2_ group (70% N_2_/30% O_2_, n=19, duration of 24h, referred to as N_2_ group), tMCAO surgery argon group (70% argon/30% O_2_, n=19, duration of 24h, referred to as argon group), tMCAO surgery or Sham surgery model construction was performed. The thrombus was removed after 3 hours of ischemia, followed by 1 hour of reperfusion. Before gas therapy, it was found that 4 mice in the tMCAO N_2_ and 4 mice tMCAO argon group each died. Autopsy showed that all the mice died from cerebral hemorrhage. Gas therapy was performed in the Attendor 120 Pro animal gas control system (Huayi Ningchuang Co., Ltd, Zhejiang Province, China).

On the 7th day after gas therapy (including the 24h of gas therapy), the experimental group and control group mice were evaluated for their neurological scores. The neurological score was based on the modified Garcia scoring system used in our previous published paper [27]. All operations in this animal experiment comply with the animal experiment standards of the Ethics Committee of Shengjing Hospital affiliated to China Medical University No. 2021PS236K(X2).

### 2.2 TTC staining measurement of cerebral infarction volume in mice

TTC staining and measurement of the cerebral infarction area were performed in mice (operate according to the instruction manual). The volume of cerebral infarction was measured and calculated by using Image Pro Plus 6.0 software. The infarct volume ratio was calculated as V1/V2×100%, where V1=∑S1×d and V2=∑S2×d, (S1: infarct area of each section; S2: total area of each section; d: thickness of each section).

### 2.3 Establishment of an oxygen glucose deprivation reperfusion model in a mouse hippocampal neuron cell line and gas therapy

We used the mouse hippocampal neuron cell line HT22 cells (purchased from the Chinese Academy of Sciences Shanghai Cell Bank) to establish the OGDR model in vitro. The HT22 cell line was cultured in a high glucose DMEM medium with 10% fetal bovine serum and passaged three times, at 37℃, and 5% CO_2_. When the cell density reached about 70%, we passaged the cells into a 10cm cell culture dish, replaced the culture medium with sugar free DMEM medium, and immediately placed the cell culture dish in a low oxygen cell culture incubator containing 1% O_2_, 5% CO_2_, and 94% N_2_ for 3h. We then changed to high glucose DMEM culture medium and incubated for 1h under normal atmospheric conditions (simulating the reperfusion process in vitro). Mirroring the different experimental groups, we placed the cell culture dishes in environments containing 70% argon, 5% CO_2_, and 25% O_2_ as the experimental group, or in an environment containing 70% N_2_, 5% CO_2_, and 25% O_2_ as the control group.

### 2.5 CCK-8 (Cell Counting Kit-8) experiment on lactate dehydrogenase (LDH) release rate

CCK-8 experiment: a HT22 cell suspension was prepared for cell counting, and seeded into a 96 well plate with approximately 100µl (2×10^3^ cells/well) per well. Each group of cells was seeded into 5 wells as replicates, and the 96 well plate was cultured in a 37℃, 5%CO_2_ incubator until the cells adhered to the wall. After OGDR modeling of H1299 cells, 10ul of CCK-8 wase was added to each well, and the plate was incubated in the dark for 1 hour. Absorbance at 450nm was measured continuously for 4 days at the same time every day, and plotted as a cell growth curve, the experiment was repeated three times.

LDH release rate experiment: A HT22 cell suspension was added to each well (100µl, 5×10^2^cells/well), with 5 additional wells per group serving as repeats. The following operation was according to the instruction manual. Absorbance at 490nm was measured, and the LDH releasing rate was calculated as follows: LDH releasing rate (%) = (absorbance of processed sample-absorbance of sample control well)/(absorbance of maximum enzyme activity of cells-absorbance of sample control well)×100%, the experiment was repeated three times.

### 2.6 Western blot

Briefly: first, extract the total protein of each group of cells, and then test the protein concentration by the BCA method. After SDS-PAGE gel electrophoresis and membrane transfer and sealing, add the first antibody and incubate at 4℃ overnight. The first antibodies included: P62 (Abcam, 1:1000, rabbit derived polyclonal antibody, ab91526), LC3 (Sigma Aldrich, 1:500, rabbit derived polyclonal antibody, ABC929), β-actin (MCE, 1:5000, mouse monoclonal antibody, HY-P80438), PI3K (Abcam, 1:1000, rabbit derived monoclonal antibody, ab302958), p-PI3K (Abcam, 1:1000, rabbit derived polyclonal antibody, ab138364), Akt (MCE, 1:2000, rabbit monoclonal antibody), p-Akt (MCE, 1:1000, rabbit monoclonal antibody, HY-P80276), mTOR (MCE, 1:1000, rabbit polyclonal antibody, HY-P80276), p-mTOR (MCE, 1:1000, rabbit monoclonal antibody, HY-P80837). Afterwards, add secondary antibody as needed and incubate at 37 ℃ for 2h, followed by ECL luminescence. The experiment was repeated three times.

### 2.7 Observation of autophagosomes by transmission electron microscopy

The cells treated with gas therapy were first added to electron microscopy fixative and fixed at room temperature in the dark for 5 minutes. The following operation was according to the instruction manual. Twenty cells were randomly selected, and the number of autophagosomes was counted. The average and standard deviation were calculated and compared.

### 2.8 Application of small molecule agonists and inducers

To verify whether argon gas therapy can regulate autophagy by modulating the PI3K/Akt/mTOR pathway, we combined inhibitors and agonists of autophagy and the PI3K/Akt/mTOR pathway in vitro experiments based on gas therapy. The specific dosage and time were determined according to the instructions and references. The small molecule agonists and inducers including: autophagy inhibitor 3 (MedChemExpress, abbreviated as INH-3, concentration 5μM, 24h)[28], autophagy inducer 3 (MedChemExpress, abbreviated as IND-3, concentration 10μM, action 24h)[29], Akt inhibitor MK2206 (MedChemExpress, concentration: 10μM, 24h)[30], mTOR inhibitor rapamycin (MedChemExpress, concentration: 20nM, 24h)[31], Akt agonist SC79 (concentration: 8µg/ml, 24h)[32], mTOR agonist MHY1845 (concentration: 5μM; 6h)[33], these reagents were all added during added at the beginning of gas therapy.

### 2.9 Statistical analysis

After confirming a normal distribution using, a t-test and Analysis of Variance (ANOVA) was used to analyze the result of in vivo and in vitro experiments. P<0.05 is considered statistically significant. The experimental data values are reported as mean±standard deviation. All statistical analyses were conducted using GraphPad Prism software.

## 3. Results

### 3.1 Argon treatment effectively improves weight recovery, neurological function, and infarct volume, but does not alter the body temperature in a mouse CIRI model

The results of 2way ANOVA showed that there were great differences in the body weight of mice among the groups (P<0.01). The body weight of Sham N_2_ group mice on the 7th day after modeling increased compared to their initial body weight (pre-modeling: 28.93±2.74g vs post modeling: 32.93±2.28g, P<0.01), and Sham argon group (pre-modeling: 28.33±2.674g vs post modeling: 31.93±2.31g, P<0.01). But no significant difference between Sham N_2_ and Sham argon group on the first or the 7th day (Figure 1A).

**Figure 1:**
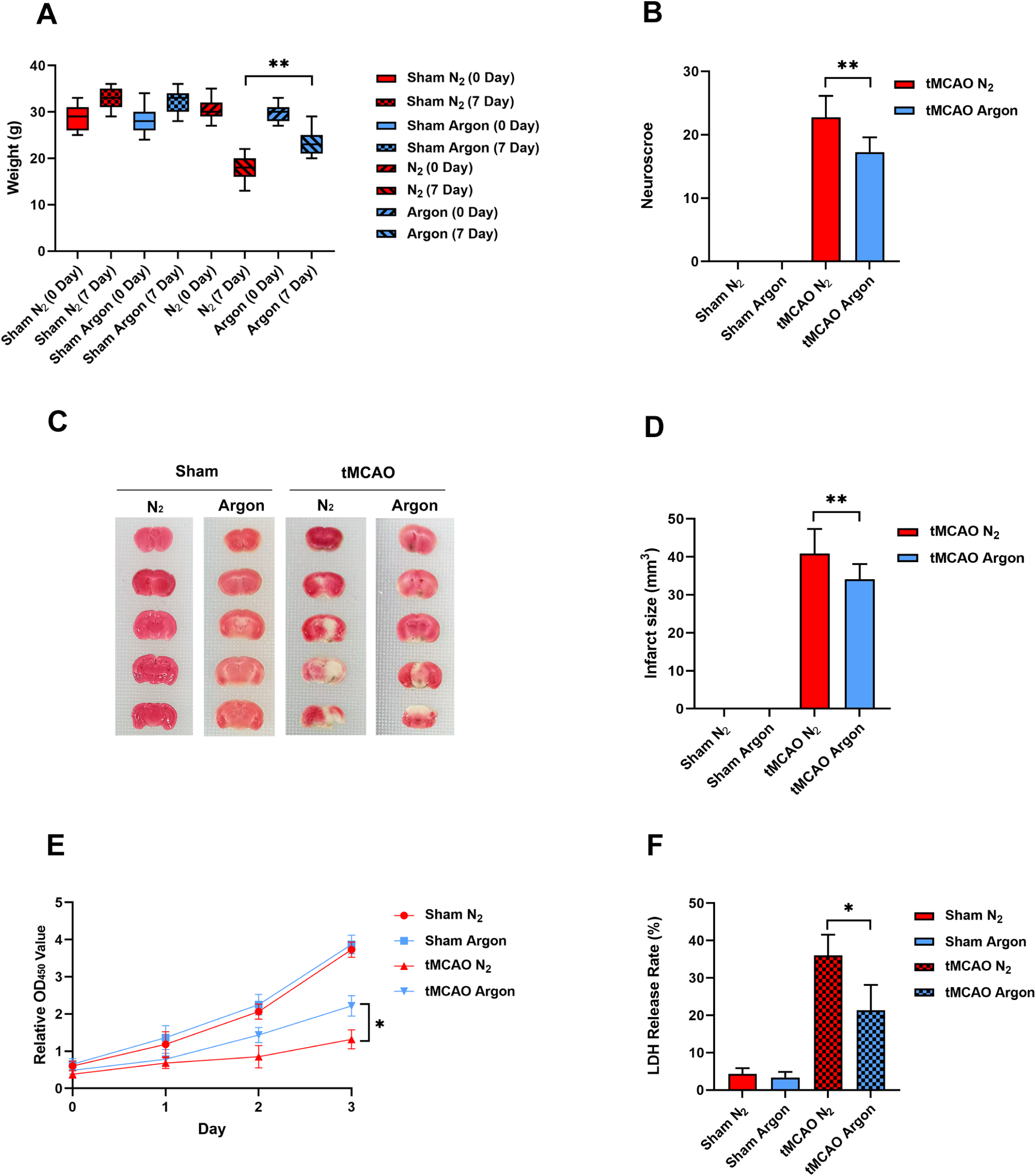
A: the average weight of mice in each group. B: the modified Garcia neurological function score of mice in the argon and N_2_ group. C: the cerebral infarction volume of mice in the argon and N_2_ group. D: bar chart corresponding to C. E: the CCK-8 results showed survival rate of HT22 OGDR model after gas treatment. F: the LDH release test of HT22 HT22 OGDR model. *: P<0.05, **: P<0.01. Sham: sham surgery group, tMCAO: transient Middle cerebral artery occlusion, Argon: argon group, N_2_: nitrogen group. Not repeated below.

Seven days after modeling and gas treatment of the mice, both the tMCAO N_2_ group and tMCAO argon group showed a decrease in average body weight (tMCAO N_2_, pre-modeling: 30.53±2.13g vs post-gas treatment: 17.87±2.45g, P<0.01; argon, pre-modeling: 29.9±1.66g vs post-gas treatment: 23.4±2.64g, P<0.01, Figure 1A). More importantly, before gas treatment, there was no significant difference between tMCAO N_2_ and agron group. However, on the 7th day after gas treatment, the average body weight of mice in the tMCAO argon group was significantly higher than that in the tMCAO N_2_ group (argon: 23.4±2.64g vs N_2_: 17.87±2.45g, P<0.01, Figure 1A).

Next, we performed a modified Garcia neurological function score on the mice on the 7th day after gas treatment. The results showed that there is no neurological damage in both Sham N_2_ and argon groups. However, the score of the tMCAO argon group was significantly lower than that of the tMCAO N_2_ group (argon: 17.27±2.31 vs N_2_: 22.73±3.41, P<0.01, Figure 1B). The TTC staining results showed that the percentage of infarct volume in the argon treatment group was lower than that in the nitrogen group (40.87±6.5% vs 34.13±3.96% respectively, P<0.01, Figure 1C and D). These results confirmed that argon treatment can improve the neurological function of the tMCAO model in mice.

The results showed the temperature before surgery were close in each group. 24h after gas therapy, there were a slightly increase in both Sham and tMCAO groups. Importantly, there was no significant difference between Sham N_2_ and Sham argon groups (37.64±0.24 VS 37.62±0.24, P>0.05); or tMCAO N_2_ and tMCAO argon groups (38.24±0.1 VS 38.13±0.1, P>0.05, Table 1).This indicates that argon treatment does not change the body temperature of mice.

### 3.2 Argon treatment significantly promotes the survival of the HT22 cell line OGDR model

CCK-8 testing results showed that there was no significant difference in the survival rate in Sham N_2_ and argon group. However, tMCAO argon group was significantly higher than the tMCAO N_2_ on the second and third day (P<0.05, Figure 1E). The results showed that the LDH release rate in the tMCAO argon group was significantly lower than that in the tMCAO N_2_ group (argon:21.33±6.8, vs N_2_: 36±5.57, Figure 1F, P<0.05), but not in Sham argon and N_2_ group. These results confirmed that argon treatment has a protective effect on the HT22 cell line OGDR model.

### 3.3 Argon treatment significantly inhibits autophagy levels

Western blot was used to detect P62 and microtubule associated protein light chain 3 (LC3) I and II. The results showed that in the Sham N_2_ and argon group, the levels of LC3 II were lower, while the levels of P62 and LC3 I were higher, and the ratio of LC3II/I was lower, which indicated a lower level of autophagy and no siginificant difference between Sham N_2_ and argon group.

After CIRI modeling, the level of LC3 II in the tMCAO N_2_ group were significantly increased compared to the Sham N_2_ group (P<0.01, Figures 2A and B), while the levels of P62 and LC3 I were significantly decreased (P<0.01, Figures 2A and B), and LC3 II/I was significantly upregulated (P<0.01, Figures 2A and B), indicating that autophagy level were significantly upregulated in tMCAO N_2_ group. Compared with the tMCAO N_2_, the level of LC3 II in tMCAO argon group was significantly downregulated (P<0.01, Figures 2A and B), and the levels of P62 and LC3 I were also significantly increased (P<0.01, Figures 2A and B). The ratio of LC3 II/I was significantly downregulated (P<0.01, Figures 2A and B). These results indicated that the autophagy levels of mice are lower under normal conditions, while mouse CIRI modeling significantly upregulates autophagy levels. Importantly, argon treatment could significantly downregulate autophagy levels in mouse CIRI models.

**Figure 2:**
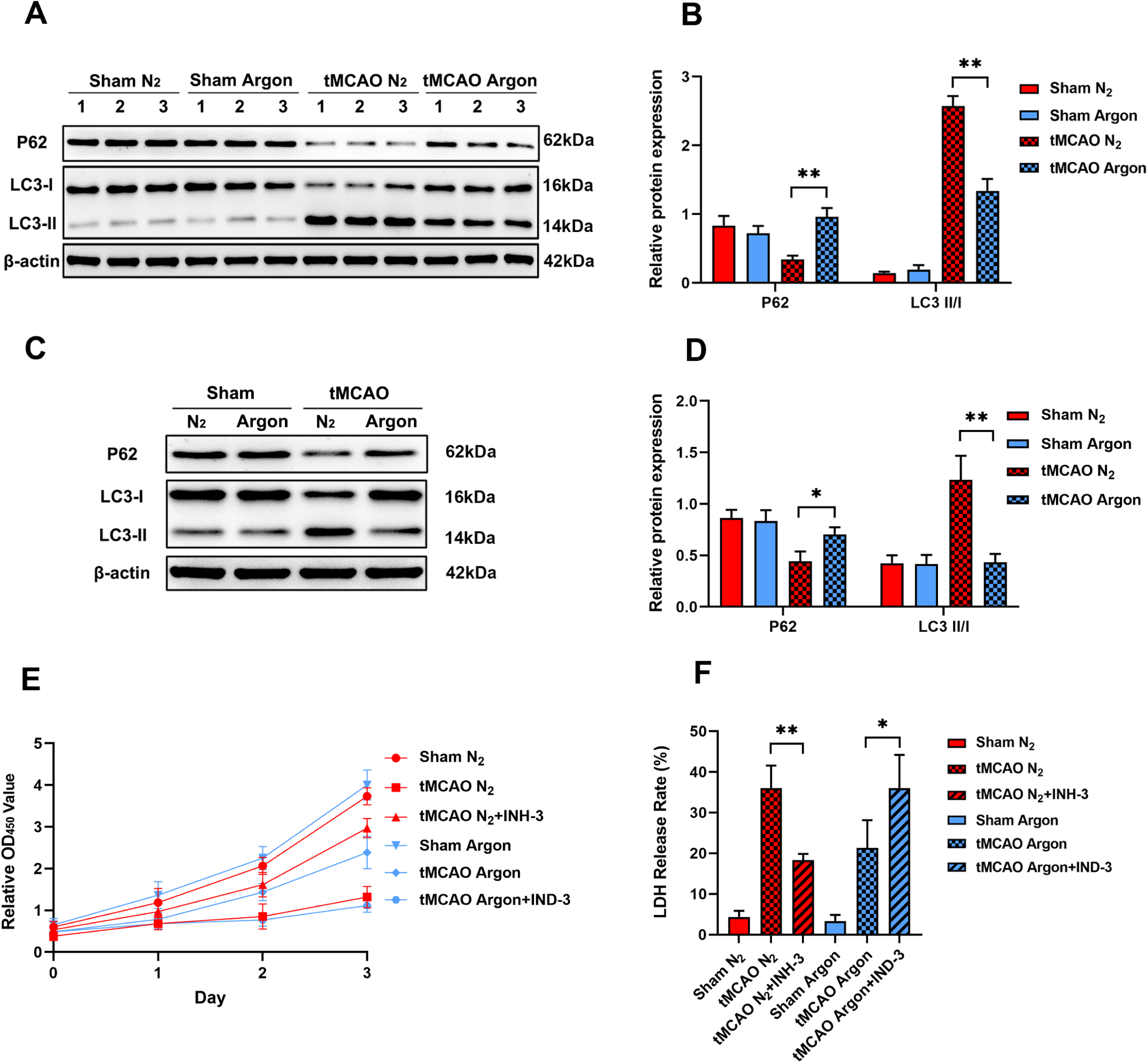
A: western blot of P62, and LC3 II/I in mice brain tissue, the Arabic numerals represent mouse numbers. B: bar chart of A. C: western blot results of autophagy proteins in N_2_ and argon groups. D: bar chart of C. E: the CCK-8 test result. F: the results of LDH release test. INH-3: autophagy inhibitor 3; IND-3: autophagy inducer 3.

The results of HT22 cells OGDR model were consistent with the results of mice model, which showed low level of autophagy in both Sham N_2_ and argon group, without significant difference (P>0.05, Figure 2C and D). Compared with the tMCAO N_2_ group, the levels of LC3 II were significantly inhibited, while the levels of P62 and LC3 I were significantly upregulated in tMCAO argon group (Figure 2C and D, P<0.05). The results indicate that argon treatment significantly inhibits autophagy levels in the OGDR model of HT22 cells.

### 3.4 Argon treatment exerts a neuroprotective function by inhibiting autophagy in HT22 cells

The CCK-8 testing results showed no significant difference of cell survial between Sham N_2_ and argon group. Compared with tMCAO N_2_ group, tMCAO combine with autophagy inhibitor 3 (INH-3) significantly promoted the survival of the HT22 cells. Compared with the tMCAO argon group, the combination of argon and autophagy inducer 3 (IND-3) significantly inhibited the survival rate of HT22-OGDR model cells (P<0.01, Figure 2E).

The results of LDH release test showed that tMCAO combine with INH-3 significantly decrease the LDH release rate in N_2_ group (P<0.01, Figure 2F). While compared with the tMCAO argon group, the combination of argon and IND-3 significantly increase the LDH release rate (P<0.05, Figure 2F).

### 3.5 Argon treatment significantly decreases autophagosomes in the HT22 cell line OGDR model

Transmission electron microscopy was used to evaluate the autophagy level of HT22 cell OGDR model after gas treatment. The number of autophagosome was counted at a magnification of 2000× and is indicated by small red arrows (Figure 3). The structure of autophagosomes was displayed at a magnification of 8000×, indicated by a large red arrow. The specific experimental groups are as follows: ①Sham N_2_; ②tMCAO N_2_; ③tMCAO N_2_+INH-3; ④Sham argon; ⑤tMCAO argon; ⑥tMCAO argon+INDH-3. The number of autophagosomes showed that there is no significant difference between Sham N_2_ and argon group. However, the number of autophagosome in the tMCAO argon group was significantly less than that in the tMCAO N_2_ group (N_2_: 7.85±1.31 vs argon: 3.25±1.02, P<0.01, Figure 3).

**Figure 3:**
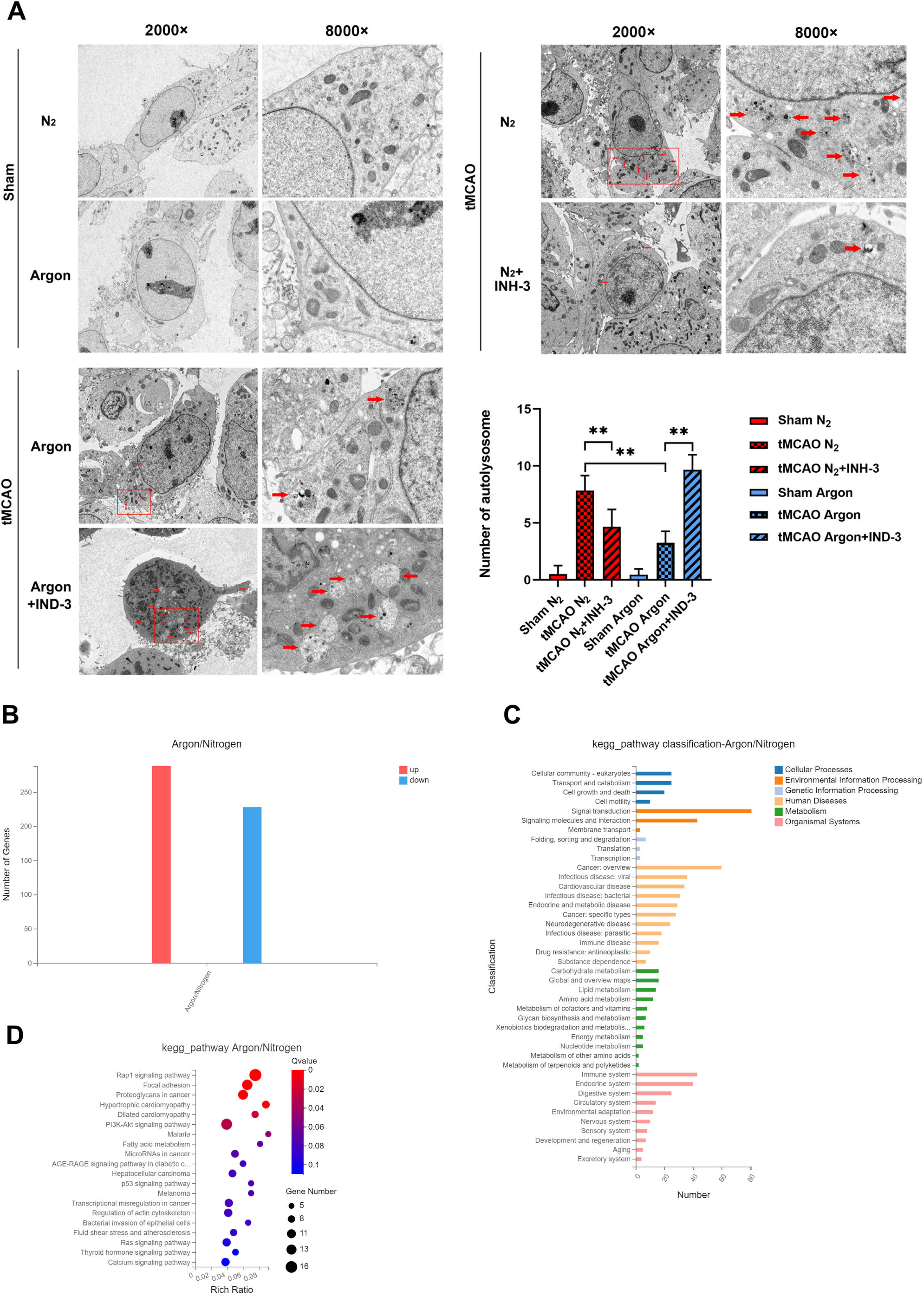
A: The number of autophagosomes was counted by transmission electron microscopy. The autophagosomes were marked with red arrows. The magnification was 2000×on the left and 8000×on the right. B: Differential gene analysis results after transcriptome sequencing. C: Classification analysis of KEGG signaling pathways, the vertical axis represents classification, and the horizontal axis represents the number of genes. D: Enrichment analysis of KEGG signaling pathway showed that the differentially expressed genes upregulated in the argon group were significantly enriched in the PI3K/Akt signaling pathway.

Transmission electron microscopy results showed that the N_2_ combined with INH-3 significantly reduced the number of autophagosomes compared to the N_2_ treatment alone (tMCAO N_2_: 7.85±1.31 vs tMCAO N_2_+INH-3: 4.65±1.531, P<0.01, Figure 3C and D). Moreover, after the combined application of IND-3, the number of autophagosomes in the argon group significantly increased (tMCAO argon: 3.25±1.02 vs tMCAO argon+IND-3: 9.65±1.35, P<0.01, Figure 3). These results indicated that argon therapy inhibits autophagy, and this effect can be reversed by an autophagy inducer.

### 3.6 Bioinformatics analysis shows that the PI3K/Akt pathway is activated in HT22 cells treated with argon gas

We collected HT22-OGDR model cells 24h after argon therapy and extracted total RNA for transcriptome sequencing and bioinformatics analysis (BGI Genomics, Dr. TOM online analysis system). Differential expression analysis showed that compared with the N_2_ group, there were a total of 516 differentially expressed genes, including 288 up-regulated genes and 228 down regulated genes (Figure 3B).

Subsequently, we classified and analyzed the differentially expressed genes through KEGG (Kyoto Encyclopedia of Genes and Genomes) signaling pathways.

The results showed that signal transduction and environmental information processing were the most active amongthe signaling pathway classifications (Figure 3C, with a total of 81 genes). Next, we performed KEGG signaling pathway analysis on differentially expressed genes, and the results showed that upregulated genes were highly enriched in the Rap1 signaling pathway, Focal adhesion, Proteoglycans in cancer, Dilated cardiomyopathy, Phosphatidylinositol 3-kinase (PI3K)/protein kinase B (PKB/Akt) signaling pathway, among others. Among them, the PI3K/Akt pathway is closely related to cell survival and death (Figure 3D).

### 3.7 Argon treatment activates the PI3K/Akt/mTOR signaling pathway and inhibits autophagy

Western blot analysis showed that the total protein levels of PI3K, Akt, and mTOR in the Sham and tMCAO group. For Sham group, there were no significant differece of p-PI3K, p-Akt and p-mTOR levels between N_2_ and Argon treatment. Compared with Sham N_2_ group, the expression of p-PI3K, p-Akt and p-mTOR in tMCAO group was significantly inhibited (P<0.01, Figures 4A and B). However, the expression of p-PI3K, p-Akt and p-mTOR tMCAO were upregulated in tMCAO argon comparing with tMCAO N_2_ group (P<0.01, Figures 4A and B), suggesting that argon can activate the PI3K/Akt/mTOR signaling pathway in the case of neurological damage.

**Figure 4:**
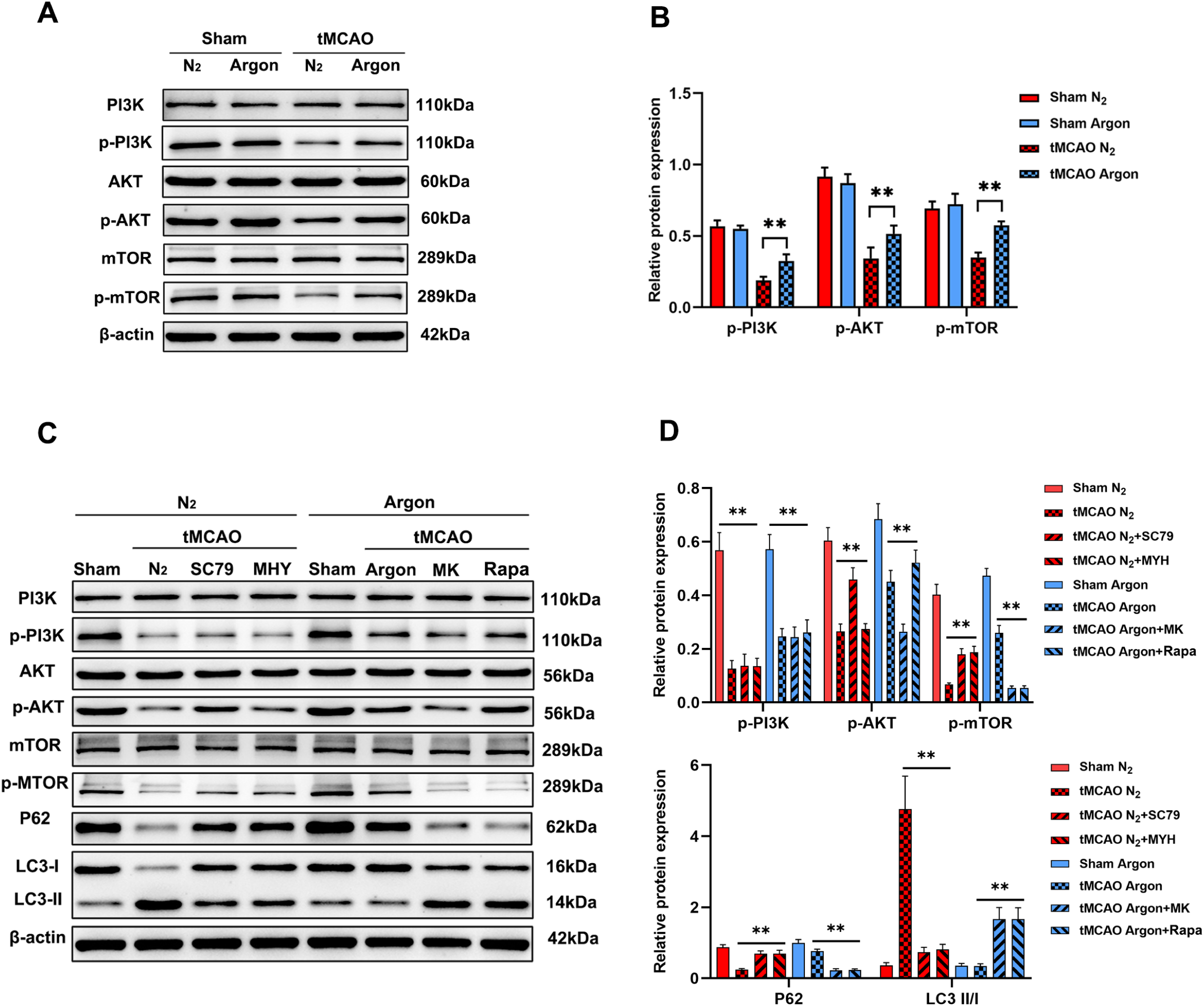
A: western blot results showed that argon treatment significantly upregulated the levels of p-PI3K, p-Akt, and p-mTOR. B: Bar chart for A; C: western blot results of PI3K/Akt/mTOR pathway factors and autophagy factors in N_2,_ N_2_ combined with SC79, and N_2_ combined with MHY1485 group; D: Bar chart for C. E: western blot results of in argon, argon combined with MK2206 and argon combined with rapamycin groups. F: Bar chart for E. MHY: MHY1485; Rapa: rapamycin.

Firstly, we added Akt agonist SC79 or mTOR agonist MHY1845 to the tMCAO N_2_ group. The expression of p-Akt was significantly upregulated (P<0.01, Figures 4C and D) after SC79 was added, and the downstream p-mTOR was also significantly upregulated (P<0.01, Figures 4C and D), accompanied by the downregulation of LC3II/I (P<0.05) and the upregulation of P62 (P<0.05, Figures 4C and D). Secondly, the combined use of MHY1845 led to significant upregulation of p-mTOR (P<0.01, Figures 4C and D), accompanied by significant downregulation of LC3II/I expression levels (P<0.01) and significant upregulation of P62 levels (P<0.01, Figures 4C and D).

Thirdly, we added Akt inhibitor MK2206 to the tMCAO argon group. The results showed that after adding MK2206, compared with the application of argon alone, the levels of p-Akt and p-mTOR were significantly decreased (P<0.01, Figure 4C and D), accompanied by a significant upregulation of LC3II/I expression levels (P<0.01) and a significant downregulation of P62 levels (P<0.01, Figure 4C and D). Fourthly, mTOR inhibitor rapamycin was added into the tMCAO argon group. The results showed that p-Akt remained unchanged, but the level of p-mTOR was significantly downregulated, accompanied by a significant upregulation of LC3II/I (P<0.01, Figure 4C and D) and significant downregulation of P62 (P<0.01, Figure 4C and D). The above results indicate that the addition of an p-Akt blockade or p-mTOR blockade can significantly block the ability of argon to inhibit autophagy. Therefore, these results confirmed that argon inhibits autophagy by activating the PI3K/Akt/mTOR pathway.

### 3.8 Argon treatment inhibits autophagy by activating the PI3K/Akt/mTOR signaling pathway

Transmission electron microscopy results showed low level of autophagy in Sham N_2_ and argon group without difference. The tMCAO nitrogen treatment group had the same experimental components as above: ①N_2_; ②N_2_+SC79; ③N_2_+MHY1485. The results showed that the number of autophagosomes in the N_2_ group was significantly higher than that in the N_2_+SC79 group and N_2_+MHY1485 group (N_2_: 8.15±1.39 vs N_2_+SC79: 3.6±1.43, P<0.05; N_2_: 8.67±1.7 vs N_2_+MHY1485: 3.25±1.07, P<0.05, Figures 5A). The tMCAO argon group is classified as above: ①argon; ②argon+MK2206; ③argon+rapamycin. The results showed that the number of autophagosomes in the argon group was significantly lower than that in the argon+MK2206 group and argon+rapamycin group (argon group: 4.05±1.32 vs argon+MK2206: 11.2±1.7, P<0.01; argon group: 4.05±1.32 vs argon+rapamycin: 10.55±1.79, P<0.01, Figure 5A).

**Figure 5:**
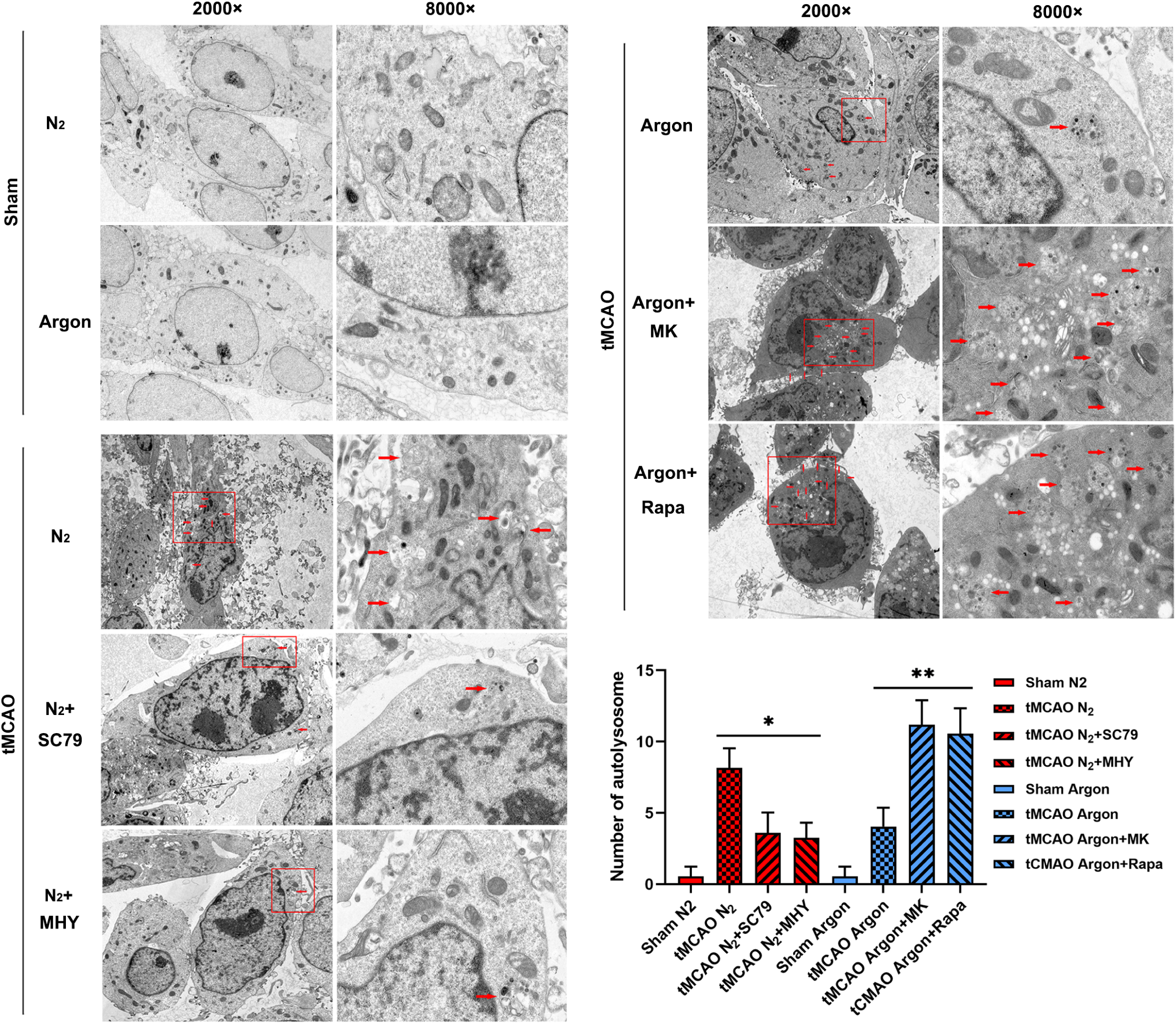
The number of autolysosomes in the tMCAO N_2_ group was significantly higher than that in the tMCAO N_2_+SC79 group, as well as the tMCAO N_2_+MHY1485 group. The number of autolysosomes in the argon group was significantly lower than that in the argon+MK2206 and argon+ rapamycin groups. Sham N_2_ and argon group serve as control.

### 3.9 Argon treatment inhibits autophagy and exerts a neuroprotective function by activating the PI3K/Akt/mTOR pathway

The CCK-8 results showed that the cell survival of the N_2_ group was significantly lower than that of the N_2_+SC79 group and N_2_+MHY1485 (P<0.05, Figure 6A), and the cell survival rate of the argon group was significantly higher than that of the argon+MK2206 and argon+rapamycin group (P<0.05, Figure 6B). The results of LDH release test showed that the LDH release rate of N_2_ group was higher than that of the N_2_+SC79 and N_2_+MHY1485 groups (P<0.05, Figure 6C), while the LDH release rate of the argon group was significantly lower than that of the argon+MK2206 and argon+rapamycin groups (P<0.05, Figure 6D). These results indicated that argon treatment inhibits autophagy and exerts a neuroprotective function by activating the PI3K/Akt/mTOR pathway.

**Figure 6:**
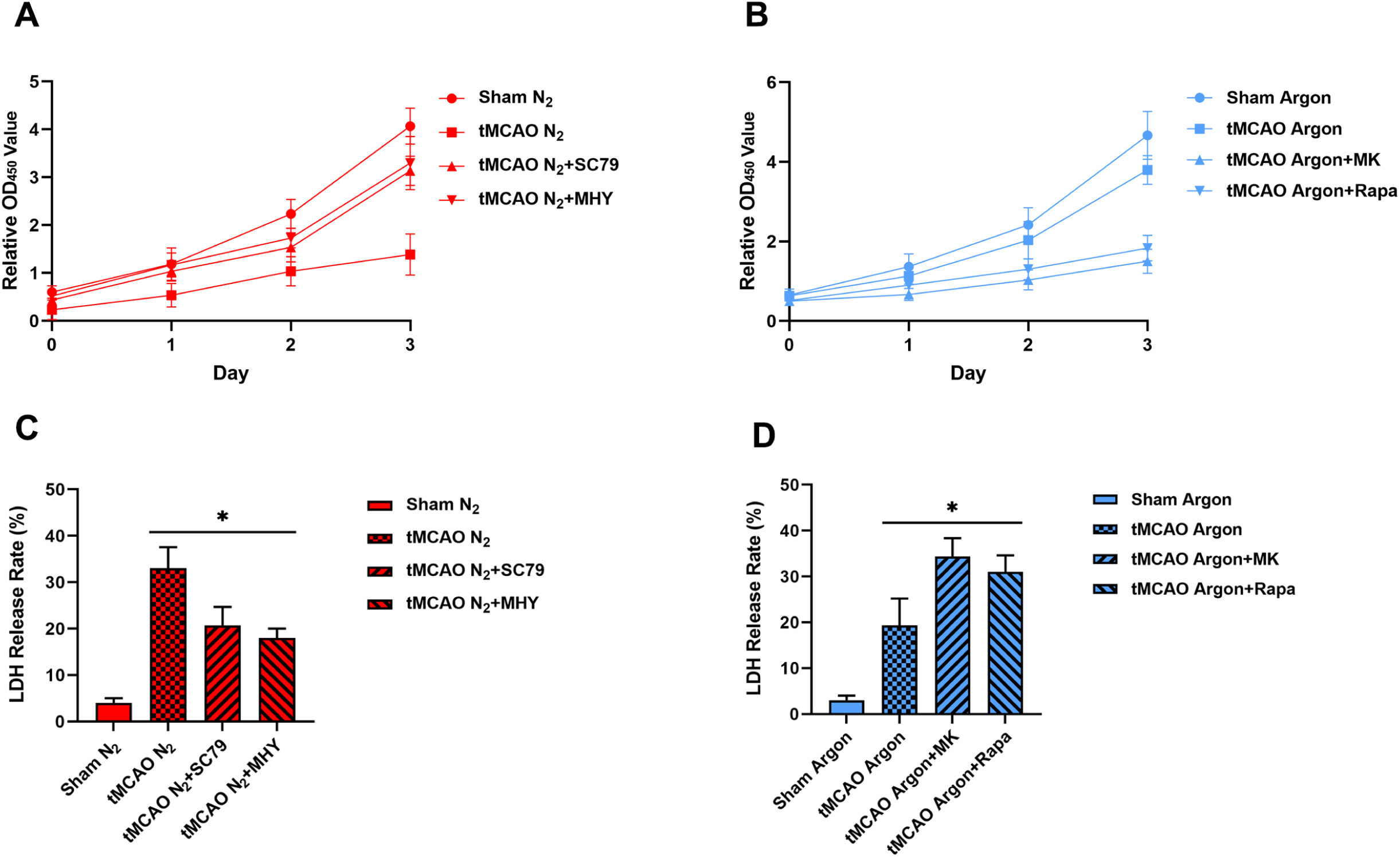
CCK-8 results show that: A: The cell survival rate of the tMCAO N_2_ group is significantly lower than that of the tMCAO N_2_+SC79 and tMCAO N_2_+MHY1485 groups. B: The cell survival rate of the tMCAO argon group was significantly higher than that of the tMCAO argon +MK2206 and tMCAO argon+rapamycin groups. The LDH release test showed that: C: The LDH release rate of the tMCAO N_2_ group was higher than that of the tMCAO N_2_+SC79 and tMCAO N_2_+MHY1485 groups; D: The LDH release rate of the argon group was significantly lower than that of the argon+MK2206 and argon+rapamycin group, *: P<0.05. Sham N_2_ and argon group serve as control.

## 4. Discussion

Our previous study showed that argon gas treatment significantly improved the neurological function scores and weight recovery of the CIRI model rats (ischemia for 90 minutes, immediate argon gas treatment for 24h after removing the plug), but did not significantly improve the infarct area[27]. We also noticed that Ryang et al. conducted a tMCAO model in rats for 120 minutes. One hour later, the animals receive argon treatment (50% argon, 50% O_2_) or nitrogen (50% N_2_, 50% O_2_) for 1h. In animals treated with argon, the infarct volume in the cortical and basal ganglia regions decreased^[14]^. Therefore, we speculate that argon gas may exhibit more significant neuroprotective effects in severe brain injuries. We applied a mouse brain CIRI model (ischemia time of 3h, reperfusion time of 1h) for argon treatment. The results showed that the neurological function, infarct size, and weight recovery of mice were significantly improved, suggesting that argon may have a more significant neuroprotective effect in severe brain injuries.

At present, research on the mechanism of argon’s neuroprotective function mainly focuses on anti-apoptosis and neuroinflammation[17, 34]. Several signaling pathways related to argon’s neuroprotective function have been identified, which including TLR2 and TLR4, HO-1 and hypoxia inducible factor-1 α (HIF-1α) and so on[35, 36]. Studies also show indicate that cerebral CIRI can activate autophagy, which not only protects cells in nutrient deficient states but also promotes cell survival. However, autophagy is a “double-edged sword,” as excessive activation of autophagy during CIRI can lead to cell lysis and promote cell death. Inhibiting this excessive autophagy may be an effective method to improve CIRI[20–26].

We first utilized transcriptional sequencing and KEGG pathway analysis in HT22 cell line OGDR model with agron treatment, and finally identified the PI3K/Akt pathway closely related to the neuroprotective function of argon gas. As is well known, the PI3K/Akt pathway is a classic autophagy signaling pathway and an upstream regulator of mTOR, playing an important role in mTOR activation. Studies have shown that transplantation of bone marrow mesenchymal stem cells can promote neurological recovery, reduce cerebral infarction volume, which may be related to activating the PI3K/Akt/mTOR signaling pathway and inhibiting excessive autophagy[37–39]. Our transcriptome sequencing results showed that the PI3K/Akt pathway was highly activated in HT22 cells treated with argon. We applied Akt agonist SC79 and mTOR agonist MHY1485 in the N_2_ group, and the results showed that both can significantly inhibit the autophagy level and promote cell survival. These results indicates that in the HT22 cells CIRI model, activating PI3K/Akt/mTOR can inhibit autophagy and exert a neuroprotective function. Similarly, in the argon group, PI3K/Akt is in an activated state. Therefore, we found that both Akt inhibitor MK2206 and mTOR inhibitor rapamycin can significantly inhibit the PI3K/Akt/mTOR pathway and block the neuroprotective function of argon treatment. These results confirm that argon activates the PI3K/Akt/mTOR pathway, inhibits autophagy, and exerts neuroprotective effects through bidirectional regulation of the PI3K/Akt/mTOR pathway. Zhao et al. found through immunofluorescence staining that argon treatment could upregulate the expression of p-mTOR and nuclear factor (erythroid derived 2)-like 2 (Nrf2) and exert neuroprotective functions. These studies supported our research results well[18].

In summary, in both in vivo and in vitro CIRI models, our results suggest that argon therapy may have significant neuroprotective effects on relatively severe nerve damage. Prolonged high concentration argon therapy may exert neuroprotective effects by activating the PI3K/Akt/mTOR signaling pathway to inhibit excessive autophagy. We are also fully aware that the specific mechanism by which argon exerts neuroprotective effects is not singular. Models under different conditions and argon treatments of different concentrations and durations may exert neuroprotective effects through different mechanisms. In the future, the application of argon gas therapy and the selection of treatment duration and concentration in clinical settings require more in-depth and detailed research.

## Supplementary Material

Refer to Web version on PubMed Central for supplementary material.

## Acknowledgments

Financial Support: This study was supported by National Natural Science Foundation of China (Grant No. 81301930 to L.-H. Yang); General project of Education Department of Liaoning Province (Grant No. L2015595 to L.-H. Yang); Key R&D Program Projects of Liaoning Province (Grant No. 2018225085 to L.-H. Yang); Natural Science Fund of Liaoning Province (Grant No. 2019JH3/10300420 and 2019-MS-374 to L.-H. Yang); Natural Science Fund of Liaoning Province (Grant No. 2020-MS-142 to S. Ma); Natural Fund Guidance Plan of Liaoning Provincial Science and Technology Department (Grant No.2019-ZD-0735 to S. Ma); 345 Talent Project of Shengjing Hospital of China Medical University (Grant No. M0364 to S. Ma); Supporting the high-quality development of science and technology funding projects in China Medical University (2023020778-JH2/202, to L.-H. Yang)

## Copyright form disclosure

All the authors have disclosed that they do not have any potential conflicts of interest.

